# Structural analysis of Sub16 sedolisin of *Trichophyton rubrum* reveals a flexible nature of its pro domain

**DOI:** 10.1101/2020.01.28.922815

**Authors:** Chitra Latka, Jagga Bikshapathi, Priyanka Aggarwal, Neel Sarovar Bhavesh, Rahul Chakraborty, Sabab Hasan Khan, Bhupesh Taneja

## Abstract

*Trichophyton rubrum* is one of the leading causes of superficial skin infections worldwide. It is a keratinolytic fungus specialized in colonization of keratinized tissue of skin, hair and nails for long periods of time. The fungus encodes a wide repertoire of secreted proteases in its genome that not only aid in nutrient acquisition but also establishment of infection on the host. The proteases are synthesized in ‘prepro’ form that requires removal of the prosegment for activation. In order to gain insights into the structural association of the pro domain with the catalytic domain, we investigate the structural features of the pro domain of the secreted sedolisin member Sub16 of *T. rubrum*. Our results show that the pro domain of Sub16 may have inherent flexibility in the absence of the associated catalytic domain which is stabilized in complex with catalytic domain. This is the first report of structural investigation on a stand-alone pro domain of sedolisin family of subtilases that will help in design of further structural studies of this protein.

## 1. Introduction

Dermatophytes are a group of keratinophilic filamentous fungi that are the most common cause of superficial mycoses (or dermatophytosis) in humans and animals worldwide [1,2,3,4]. Dermatophytes can cause superficial skin infections in immunocompetent individuals while deeper skin layers might be involved in case of immunocompromised hosts [5]. Dermatophyte infections are chronic infections that may require long treatment regimes and are often associated with considerable morbidity and socio-economic trauma. Among dermatophytes, *Trichophyton rubrum* is the most common etiological agent of dermatophytosis, accounting for 70-90% of all reported dermatophytosis cases in humans [6,7,8,9]. Proteases are thought to be key factors in infection by dermatophytes as these fungi are well adapted to utilize keratin as a source of nutrient in order to infect skin, hair and nails [10].

Whole genome sequencing and comparative sequence analysis reveals enrichment of several classes of endo-and exo-peptidases in *T. rubrum* genomes [11,12,13]. Secreted proteases are required not only for nutrient acquisition from the keratinized tissues but also suggested to be important virulence factors [14,15]. Indeed, secreted proteases constitute one of the major group of enzymes in the predicted secretome of *T. rubrum* IGIB-SBL-CI1, with the MEROPS SB superfamily of subtilases [16,17] comprising of 16 members, as the most abundant secreted family [12]. Among the secreted subtilases of *T. rubrum*, 13 members belong to the S8 family (12 subtilisin-like S8A members and one kexin-like S8B member) while three members belong to the S53 (or sedolisin) family in *T. rubrum* IGIB-SBL-CI1.

Sedolisins are pepstatin-insensitive serine carboxyl peptidases that are considerably larger than subtilisins and are active at low pH [18,19]. Sedolisins, like subtilisins of S8 family, have a homologous serine residue in its catalytic triad; however, they differ from the canonical subtilases in having a conserved Ser-Glu-Asp triad instead of the Ser-His-Asp catalytic triad, characteristic of the S8 subtilisins [20]. Sedolisins are synthesized as pre-pro peptidases where the prosegment appears to be an independent domain that helps in proper folding of catalytic domain [21,22]. The ‘pro domain’ is removed via an autocatalytic intramolecular mechanism in low pH conditions [22]. Real time kinetics on human tripeptidyl peptidase 1 (TPP1), a sedolisin member, shows that its prosegment is a slow-binding inhibitor of its cognate enzyme [21]. Prosegment of human TPP1 bound to its catalytic domain [23,16] remains the only available structure of a sedolisin prosegment, while the structural details of a stand-alone pro domain from the sedolisin family are not available. The structure of precursor form of human TPP1 reveals a stable folded structure with an antiparallel-α/antiparallel-β fold of the pro domain that is packed against the catalytic domain of the protein [23,24]. In addition to their role as inhibitors of the catalytic domain, pro domains have also been ascribed the role of chaperones, aiding in correct folding and/or stability of the cognate catalytic domain both *in vivo* as well as during *in vitro* refolding experiments [25,26,27]. The significance of a functional pro domain in aiding the correct functional state of the mature protein is emphasized by identification of missense mutations in pro segment of human TPP1 associated with CLN2 disease (https://www.ncbi.nlm.nih.gov/clinvar/?term=cln2; https://www.ucl.ac.uk/ncl-disease/ncl-resource-gateway-batten-disease/mutation-and-patient-database/mutation-and-patient-datasheets-1).

In this study, we report structural features of an independent pro domain of Sub16 (NCBI Accession no. KMQ48787, Locus_tag: HL42_0530), a secreted sedolisin of *T. rubrum* through CD, NMR, preliminary crystallization and MD simulations studies in order to gain insights into the molecular structure and functions of this inhibitor domain. Our work reveals a flexible nature of the pro domain of Sub16 sedolisin in the absence of its cognate catalytic partner. We identify a short subdomain as a potential minimalistic stable component which may aid in design of potential inhibitors for sedolisins.

## 2. Material and Methods

### 2.1 Sequence analysis

Sequence homologs of Sub16 sedolisin of *T. rubrum* (NCBI accession no.: KMQ48787, Locus_tag: HL42_0530) were identified in selected filamentous fungi belonging to phylum Ascomycota, as bidirectional best hits from Basic Local Alignment Search Tool (BLAST) [28]. Multiple sequence alignment of Sub16 sedolisin sequences was carried out using Clustal Omega [29] and a phylogenetic tree was constructed using MEGA7 [30] with 1000 cycles of bootstrap.

### 2.2 Cloning and purification of Sub16 pro

*T. rubrum* IGIB-SBL-CI1 was grown for 10 days on a 0.2% soybean flour media (with soy protein as a sole carbon and nitrogen source), pH 7.0 at 30 °C. The fungal mat was then collected and homogenized in liquid nitrogen and total RNA was extracted using trizol reagent (Invitrogen, USA) followed by phase separation with chloroform and subsequent precipitation of RNA using isopropanol. DNase treatment was done to remove presence of any contaminating DNA from RNA samples. cDNA was prepared from 1 μg RNA, using Superscript cDNA synthesis kit (Thermo Fisher Scientific) following manufacturer’s instructions. The pro domain of Sub16 of *T. rubrum* IGIB-SBL-CI1, corresponding to residues 17-176 (Sub16pro), was amplified using cDNA as the template with the help of forward and reverse primers listed in Table 1. The amplified product was digested with BamHI and XhoI and cloned in similarly digested pET28-His_10_-Smt3 vector to generate pSub16. pSub16 hence encodes pro domain of Sub16 in fusion with a N-term His_10_-Smt3 tag.

**Table 1:**
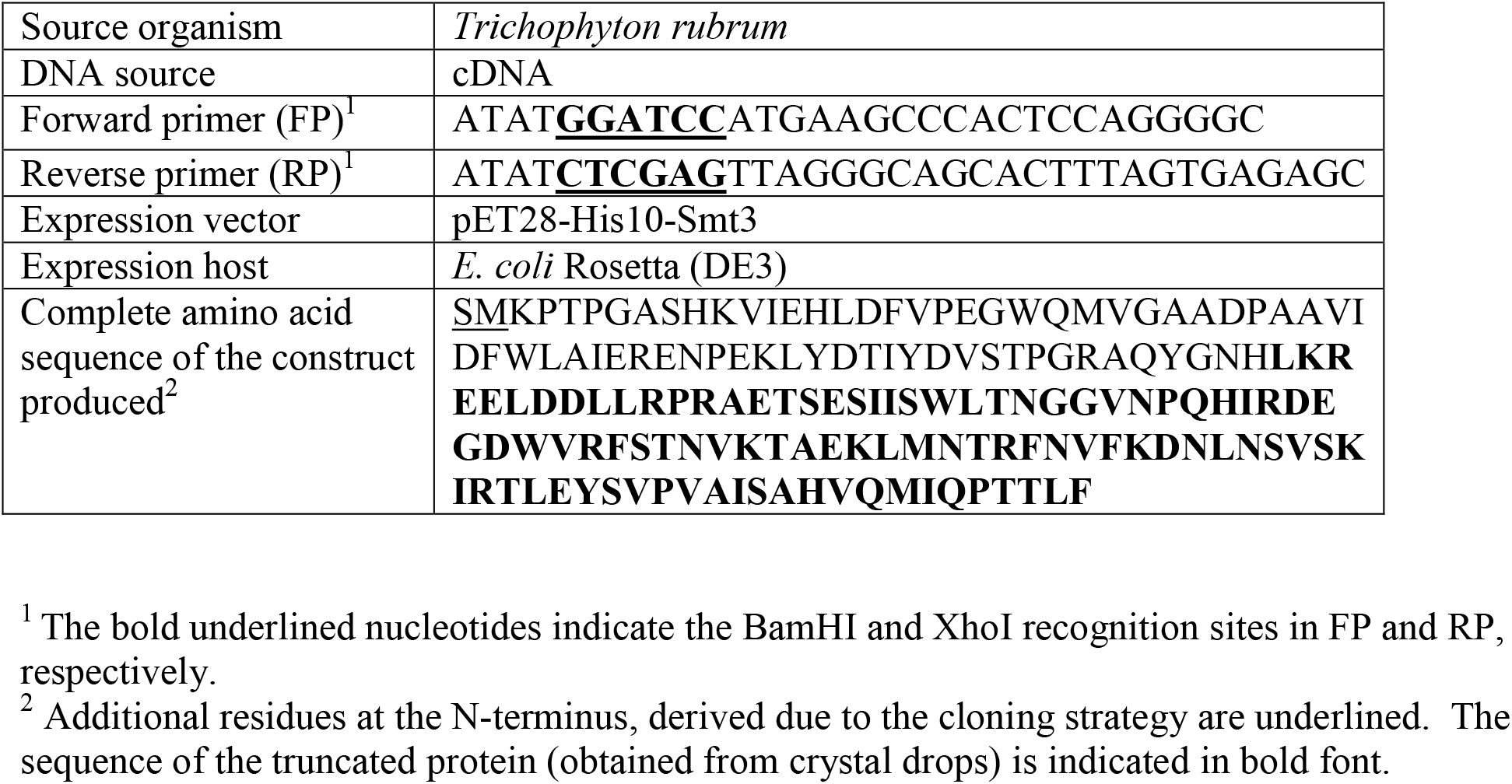
Macromolecule production information

For protein expression and purification, *E. coli* Rosetta (DE3) cells were transformed with pSub16. The tagged protein was expressed by growing the transformed cells in LB media containing 50 μg/ml kanamycin and 25 μg/ml chloramphenicol at 37 °C with constant shaking until A_600_ was 0.5. Protein expression was induced by addition of 0.3 mM isopropyl 1-thio-D-galactopyranoside. The cells were further grown with constant shaking at 16 °C for 18 h and harvested by centrifugation. All subsequent procedures were performed at 4 °C. The cell pellet was resuspended in five times its wet weight of pre-chilled lysis buffer S (25 mM Tris-HCl pH 8.0, 300 mM NaCl, 5% glycerol) containing 1mM PMSF and 1X protease inhibitor cocktail (Sigma) and lysed by sonication. The cell debris was removed by centrifugation at 16000g for 90 min and the supernatant was loaded onto a nickel-nitrilotriacetic acid-agarose (Ni-NTA) column (Thermo-Scientific) pre-equilibrated with buffer S. The column was washed thoroughly with Wash Buffer (25 mM Tris-HCl, pH 8.0, 300 mM NaCl, 5% Glycerol and 40 mM imidazole) and the bound protein was eluted with buffer S containing 300 mM imidazole.

The eluted protein was mixed with Smt3-specific protease Ulp1 (Ulp1: protein ratio as 1:500) and incubated at 4 °C overnight to cleave the His_10_-Smt3 tag. Sub16pro was separated from the cleaved tag by loading the protein on a Ni-NTA column and finally purified by size-exclusion chromatography over a Superdex 75 10/300 GL gel filtration column (GE Healthcare) and stored in buffer P (25 mM Tris-HCl, pH 8.0, 100 mM NaCl and 5% glycerol).

Molecular weight of the purified Sub16 pro was determined by spotting 1 μl of the purified sample with the matrix (sinapic acid) and analyzed on MALDI-TOF/TOF 5800 (AB Sciex) using linear mode. The MS spectra were acquired in the mass range 8 kDa to 80 kDa with laser intensity of 5500.

### 2.3 Circular Dichroism and NMR Spectroscopy

For far-UV CD spectrum measurement of Sub16pro, 6 μM protein in 10 mM sodium phosphate, pH 7.4 and 100 mM NaCl was used in a 1 mm path length quartz cuvette using a Jasco J-815 CD spectrometer equipped with Peltier-type temperature controller. CD spectrum measurement was carried out in the wavelength range 195-250 nm at 1 nm band width at 25 °C. Each spectrum was recorded as an average of three scans. Contribution of the buffer was subtracted from the spectrum of the protein and data was plotted as mean residue ellipticity (deg cm^2^ dmol^-1^).

To measure 1D NMR spectrum of Sub16pro on Bruker *Avance* III equipped with a 5 mm cryogenic triple resonance TCI probe, operating at field strength of 500.15 MHz, D_2_O was added at the final concentration of 10% (*v/v*) to the protein sample. For 2D and 3D NMR, *E. coli* Rosetta (DE3) cells were transformed with pSub16 and grown in M9 minimal media with ^13^C_6_-D-glucose and ^15^NH_4_Cl as sole source of carbon and nitrogen till A_600_ reached 0.5. Induction of protein expression and purification was carried out as described for the native (unlabeled) Sub16pro. For data acquisition, D_2_O was added at the final concentration of 10% to the protein sample and 200 μL of this was added to a 3 mm NMR tube. All experiments were measured on Bruker *Avance* III equipped with a 5 mm cryogenic triple resonance TCI probe, operating at the field strength of 500.15 MHz, at 24 °C. All NMR spectra were referenced to DSS, processed with Topspin3.1 (Bruker) and analyzed using CARA [31].

### 2.4 Crystallization, data collection and processing

Purified Sub16pro (16 mg/ml) in buffer P was used for initial crystallization screenings by mixing 0.2 μl Sub16pro with equal volume of reservoir solution by the sitting drop vapor diffusion method with the help of Mosquito crystallization robotics system (TTP Labtech) using commercially available crystallization screens (Molecular Dimensions) at three different temperatures (4 °C, 10 °C and 24 °C). Initial crystals were obtained for Sub16pro in 100 mM HEPES pH 7.5, 2 M ammonium sulfate and 2% PEG 400 at 24 °C and confirmed as protein crystals by addition of Izit dye (Hampton Research). In order to obtain crystals reproducibly, microseeding method was used with addition of 0.5 μl of seeds (1:100 dilution) in the crystallization drop.

Crystals of Sub16pro were cryoprotected in a solution containing 20% ethylene glycol in addition to the crystallization buffer (100 mM HEPES pH 7.0, 2 M ammonium sulfate and 4% PEG 400) and flash-cooled in liquid nitrogen prior to data collection. Diffraction data were collected at ID23 or ID-29 beamlines at European Synchrotron Radiation Facility (ESRF). Data was processed and integrated using iMOSFLM [32] and then scaled using SCALA of CCP4 suite [33,34]. Details of data collection and processing are provided in Table 2.

**Table 2:** Data collection and processing statistics^1^

|  |  |
| --- | --- |
| Diffraction source | ID29 ESRF |
| Wavelength (Å) | 0.97895 |
| Crystal-to-detector distance (mm) | 320 |
| Rotation range per image (°) | 0.1 |
| Total rotation range (°) | 360 |
| Exposure time per image (s) | 0.037 |
| Space group | C 222<sub>1</sub> |
| *a*, *b*, *c* (Å) | 66.72, 71.47, 72.91 |
| Resolution range (Å) | 48.77-1.68 (1.71 -1.68) |
| Total No. of reflections | 226721 (11967) |
| Total No. of unique reflections | 19993 (997) |
| Completeness (%) | 99.8 (99.7 ) |
| Multiplicity | 11.3 (12.0 ) |
| $\langle I/\sigma(I) \rangle$ | 14.3 (3.8) |
| $R_{meas}$ | 0.093 (0.929) |
| CC<sub>1/2</sub> | 0.629 |
| Wilson B factor (Å<sup>2</sup>) | 22.54 |

<sup>1</sup>Values in parentheses are for the outer shell

### 2.5 N-terminal sequencing and Mass spectrometry of Sub16 pro crystals

N-terminal sequencing of Sub16pro, harvested from crystals, was carried out by Edman degradation method [35,36]. Crystals were pooled from 5-6 crystallization drops and washed several times with crystallization buffer to remove any residual uncrystallized protein in the drop. Finally, the crystals were washed with and the protein resuspended in 100 mM HEPES pH 7.0, prior to loading on a 15% polyacrylamide gel and separation by SDS-PAGE electrophoresis. The protein was then transferred to PVDF membrane and N-terminal sequencing was carried out.

For mass spectrometry (MS) analysis of Sub16pro present in crystals, the protein harvested from the crystals was washed, resuspended in 100 mM HEPES pH 7.0 and 1 μl of the sample was spotted with matrix (sinapic acid) and analyzed on MALDI-TOF/TOF 5800 (AB Sciex) as described for the purified protein.

### 2.6 Homology Modeling of Sub16pro in apo form and in complex with catalytic domain

Homology models of Sub16pro and Sub16 catalytic domain were generated using the coordinates of human TPP1 (PDBID: 3EE6) as template [23] from MODWEB server [37].

MD Simulations were performed on the homology models of Sub16pro and Sub16 catalytic domain using GROMACS package, version 5.1.4 [38] and the OPLS-all atom forcefield [39]. The proteins were centered in a cubic box and were solvated by spc (simple point charge) of water molecules [40]. In order to obtain a neutral simulation box, respective counterions were added to Sub16 pro (6 Na^+^) or Sub16 catalytic domain (2 Cl^−^). Energy minimization of solvated systems was carried out using steepest descent followed by a conjugate gradient method until the maximum forces were smaller than 500 kJ. In all simulations, during equilibration dynamics, in order to maintain the systems in a stable environment, temperature was set to 300K by the modified Berendsen thermostat [41] with τ_T_ = 0.1 ps and pressure was set to 1 bar by the Parrinello-Rahaman algorithm [42] with τ_P_ = 2 ps. A 1.4 nm cut off was kept for short range electrostatic interactions and a particle mesh Ewald (PME) algorithm [43] was used for long range electrostatic interactions. LINCS algorithm [44] was used to constraint bond lengths with 2 fs time steps. The systems were equilibrated to 100 ps in NPT and NVT ensembles, respectively. Finally, 100 ns molecular dynamic simulation production runs were carried out at remote CSIR-4PI server.

A complex of Sub16pro with Sub16 catalytic domain was obtained from the PatchDock server [45]. Ten best solutions based on docking score and conformations were examined and a model of Sub16pro in complex with the cognate catalytic domain was selected as the starting structure for carrying out MD simulations. MD simulations were carried out for the complex as described for apo protein.

## 3. Results and Discussion

### 3.1 Sequence analysis of Sub16 sedolisin of *T. rubrum*

Whole genome sequencing of *T. rubrum* IGIB-SBL-CI1 had previously identified three members of the sedolisin/ S53 family in the secreted subset of proteases in the fungal genome [12], that were annotated as Sub14 (NCBI accession no.: KMQ45572, Locus_tag: HL42_3675), Sub15 (NCBI accession no.: KMQ42440, Locus_tag: HL42_6852) and Sub16 (NCBI accession no.: KMQ48787, Locus_tag: HL42_0530). Sub16 comprises of 596 residues with a predicted Mw of 64.8 kDa. A sequence alignment of Sub16 of *T. rubrum* with identified sedolisins of filamentous fungi belonging to phylum Ascomycota reveals several conserved sequence features (Fig. S1A), suggesting that these residues play important roles in the sedolisin family. For instance, the Glu-Asp-Ser catalytic triad, characteristic of sedolisins is conserved in all the examined fungal sedolisin sequences. A phylogenetic tree based on sedolisin sequences shows that the skin pathogens cluster together (Fig. S1B). Further examination of the sedolisin sequences of skin fungi helps identify domain boundaries in Sub16 (Fig. 1).

**Figure 1:**
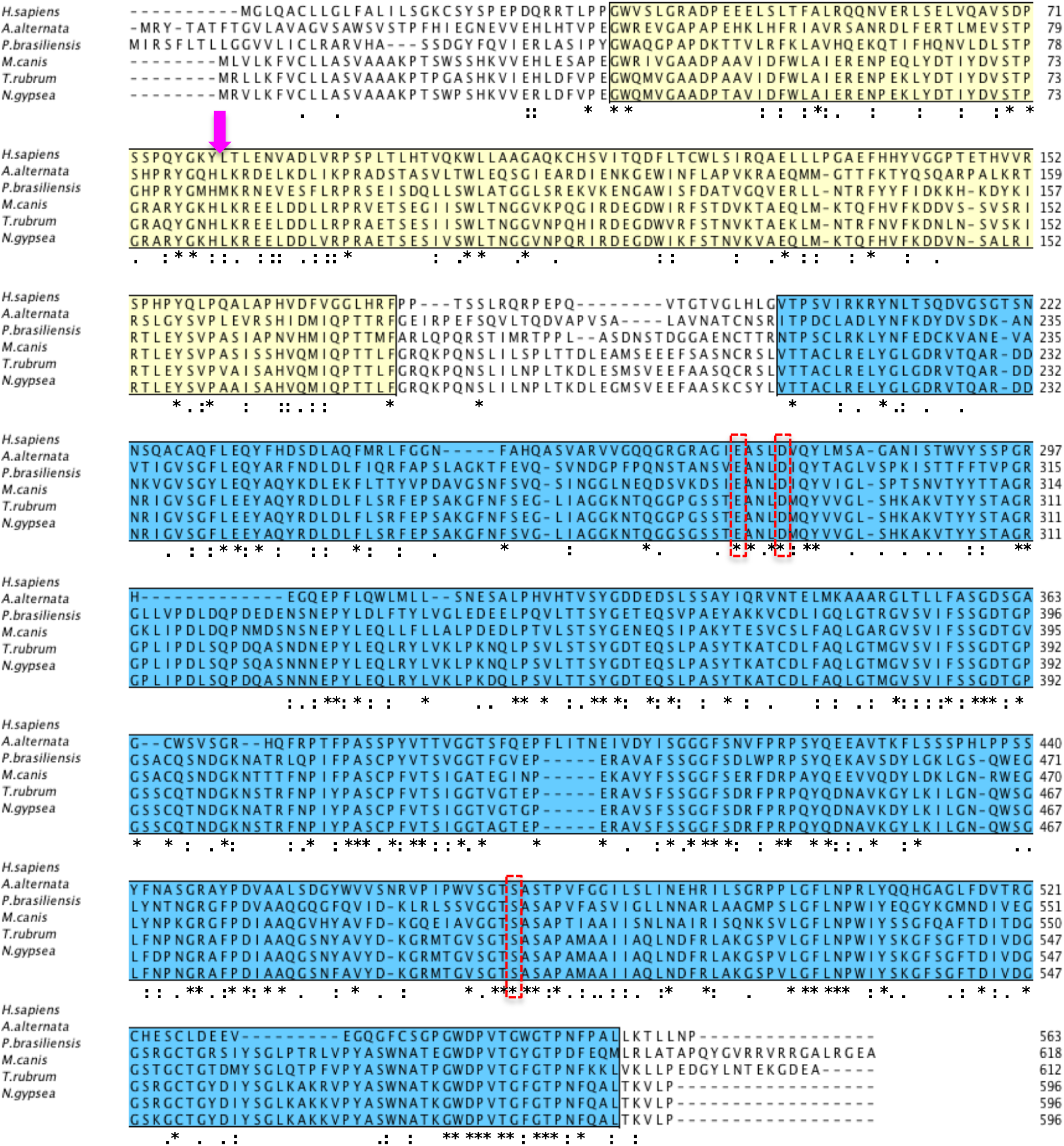
Sequence alignment of Sub16 (NCBI accession no.: KMQ48787, Locus_tag: HL42_0530) with selected sedolisin sequences of filamentous skin fungi. Pro domain is marked in yellow and catalytic domain is marked in blue. Catalytic triad is highlighted in a red box. Cleavage site of Sub16 pro domain is indicated by magenta arrow. Amino acid sequence accession numbers: *[Alternaria alternata* (NCBI accession no.: XP_018388492.1), *Microsporum canis* (NCBI accession no.: XP_002850080.1), *Paracoccidioides brasiliensis* (NCBI accession no.: ODH48645.1), *Nannizzia gypsea* (NCBI accession no.: XP_003176669.1), *Homo sapiens* (NCBI accession no.: NP_000382.3) and *Trichophyton rubrum* (NCBI accession no.: KMQ48787)].

Sub16 comprises a short variable region at the N-terminus (residues 1-16, as per Sub16 numbering) that is predicted as the signal peptide by SignalP server [46]. A conserved (^34^P-X-G/S/T/N-^37^W) and ^176^F mark the start and end of the pro domain, respectively, separated by a short linker region from the conserved catalytic domain (residues 211-596) (Fig. 1). A computational prediction to identify any potential disordered regions in the protein using DISOPRED3 [47] did not indicate any regions in Sub16pro likely to be disordered, and suggesting the protein to have an ordered α/β structure, as also indicated by secondary structure prediction of Sub16pro by GOR4 [48] or CFSSP [49] (not shown).

### 3.2 Purification of Sub16pro

Sub16pro was cloned in and successfully overexpressed from pET28-His_10_-Smt3 vector. The protein was purified to homogeneity from soluble portion of *E. coli* lysate over a Ni-NTA resin followed by gel filtration chromatography (Fig. S2). The purified protein had a deduced molecular weight of 18.1 kDa based on its amino acid sequence, which is in agreement with the experimentally determined molecular weight of 18.3 kDa by mass spectrometry.

In order to determine any potential subdomain movements in Sub16pro due to the absence of the bound catalytic domain, a 1D ^1^H NMR spectrum of Sub16pro in solution (10 mM sodium phosphate pH 7.4 and 100 mM NaCl) was acquired on a Bruker AvanceIII spectrometer operating at 500.15 MHz. A1D ^1^H NMR spectrum of Sub16pro along with the CD spectrum is indicative of a folded protein with a mixed α/β structure (Fig. 2A and B).

**Figure 2:**
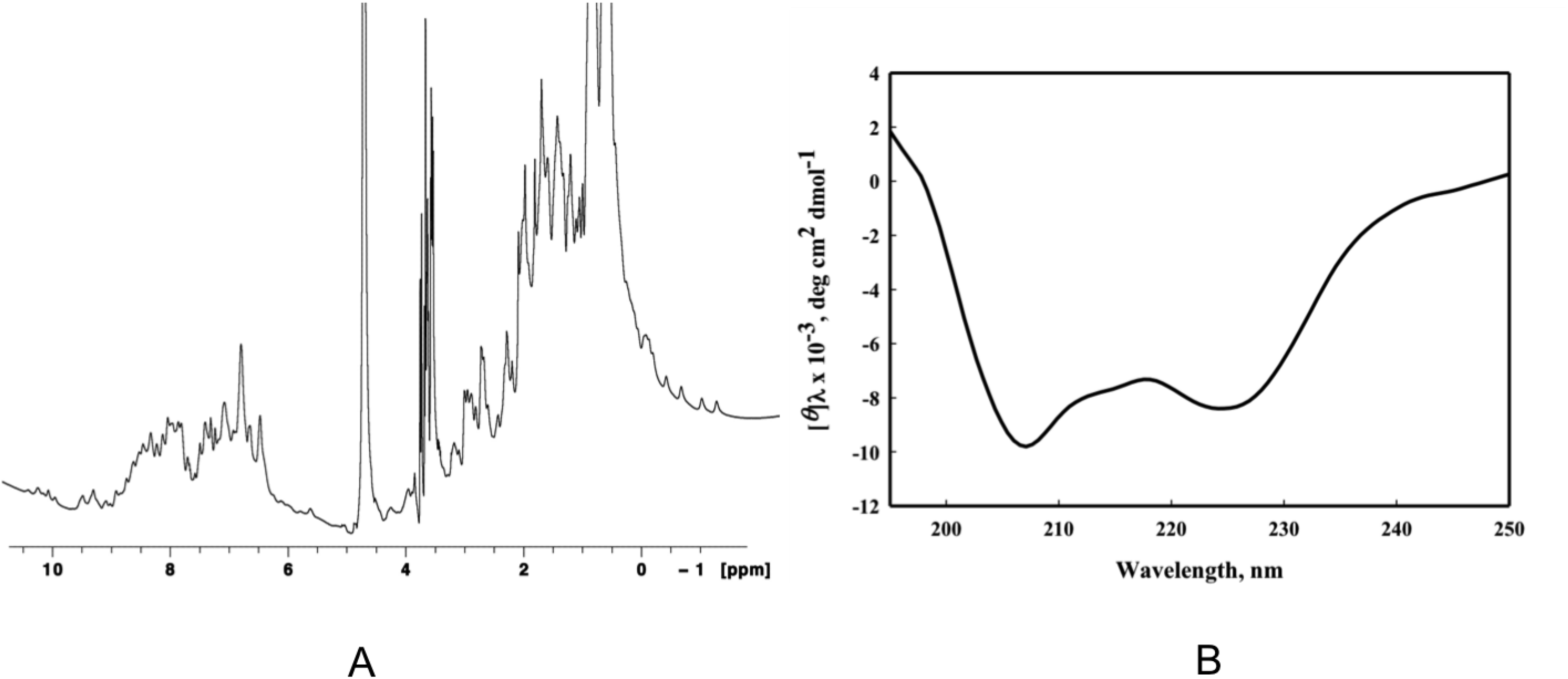
Structural analysis of Sub16pro. (A) 1D NMR spectrum of Sub16pro and (B) CD analysis of Sub16pro. Both spectra are characteristic of folded proteins.

### 3.3 Crystallization and identification of a minimalistic subdomain

Protein crystallization can serve as a useful tool to assess the purity as well as structural homogeneity of a protein molecule. In order to obtain further insights into the structural homogeneity of recombinant Sub16pro and carry out structural investigations, crystallization was set up with the purified protein. Initial crystallization experiments resulted in small rod-shaped crystals in 100 mM HEPES, pH 7.5, 2 M ammonium sulfate and 2 % PEG 400 as a precipitant. Diffraction-quality crystals, however, could be produced only upon inclusion of ‘crushed’ initial crystals as crystallization nuclei in the crystallization drops by microseeding method against a reservoir solution of 100 mM HEPES pH 7.0, 2 M ammonium sulfate and 4 % PEG 400. Several diffraction datasets were collected with these crystals on the ID23 and ID29 beamlines (ESRF). The highest quality X-ray diffraction data set was collected at ID29 to a resolution of 1.68 Å suggesting protein in the crystal exists in a folded form. A summary of data collection and processing statistics is given in Table 2. Sub16pro shares low sequence similarity (< 30%) with prosegment of human TPP1. Attempts to solve the structure by molecular replacement using the pro domain of human TPP1 (PDB IDs: 3EE6 [24], 3EDY [23]) or a homology model of Sub16pro as search models were, however, surprisingly, unsuccessful.

The absence of a solution by molecular replacement and the lack of reproducibility without microseeding persuaded us to further investigate the nature of Sub16pro in the crystals. Sub16pro crystals were harvested from crystallization drops and the protein analyzed on a 18% polyacrylamide gel by SDS-PAGE electrophoresis. Only a ~11 kDa band was observed for the protein harvested from crystals, instead of the intact ~18kDa band of the native protein by SDS-PAGE (Fig. 3A). MS analysis of the two bands indicated sizes of 18.3 kDa and 11.1 kDa, corresponding to sizes of native full-length Sub16pro or a truncated fragment, respectively (Fig. 3B & C). Finally, N-terminal sequencing of the truncated protein obtained from the crystals (Fig. 3A, lane 2) was carried out by Edman degradation method. N-terminal sequencing revealed an amino acid sequence of LKREELDD, indicating that the protein in the crystal is composed of residues 82-176 of Sub16 (Fig. 1, Table 1), with a deduced molecular weight of 10.8 kDa based on its amino acid sequence which is in excellent agreement with the 11.1 kDa as determined by mass spectrometry, and is likely the most stable form of the pro domain that yielded diffraction-quality crystals. A molecular replacement solution for the truncated protein, with shorter sub-structures of human TPP1 as search models, however, did not result in a solution.

**Figure 3:**
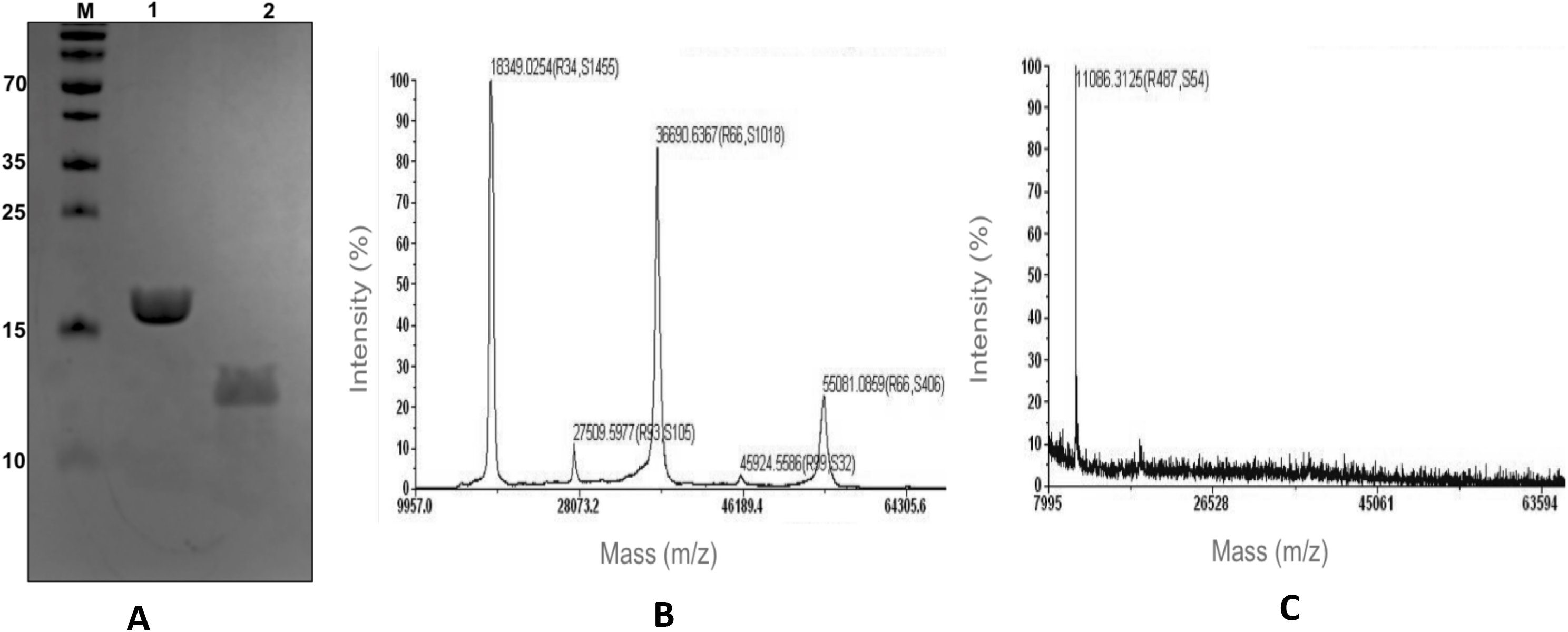
Analysis of Sub16pro present in crystallization drops. (A) SDS-PAGE profile of Sub16pro. Lane 1: Native purified Sub16pro; lane 2: Sub16pro harvested from crystal drops. Lane M indicates molecular weight standards (in kDa). Mass spectrometry profile of (B) native full-length Sub16pro (before crystallization). or (C) Sub16pro harvested from crystal drops.

### 3.4 NMR spectroscopy studies

In the absence of protein crystals for the complete unit of Sub16pro (corresponding to residues 17-176 of Sub16), we decided to investigate the three-dimensional structure of the standalone Sub16pro by NMR spectroscopy through acquisition of 2D and 3D NMR spectra like 2D [^15^N, ^1^H] HSQC, 2D [^13^C, ^1^H] HSQC, 3D HNCA, 3D HNCACB, 3D CB(CA)CONH, 3D HNCO, 3D HN(CA)CO spectra on double labeled ^13^C and ^15^N labeled protein. However, the data that was obtained for Sub16pro suggested that Sub16pro is forming higher order structures and a possible conformation exchange at the μs-ms time scale (Fig. 4), limiting further use of this method. This suggests that Sub16pro has some inherent flexibility that might be stabilized by the catalytic domain *in vivo*. In the absence of the cognate catalytic domain, the pro domain forms a higher order structure, as determined by NMR or results in a truncated fragment (in crystallization drops), thus, posing hurdles in its three-dimensional structure determination by these experimental methods. The fragment comprising residues 82-176, identified in the protein crystals, appears to be the minimalistic stable subdomain form of Sub16pro. However, production of this subdomain from a recombinant clone was unsuccessful.

**Figure 4:**
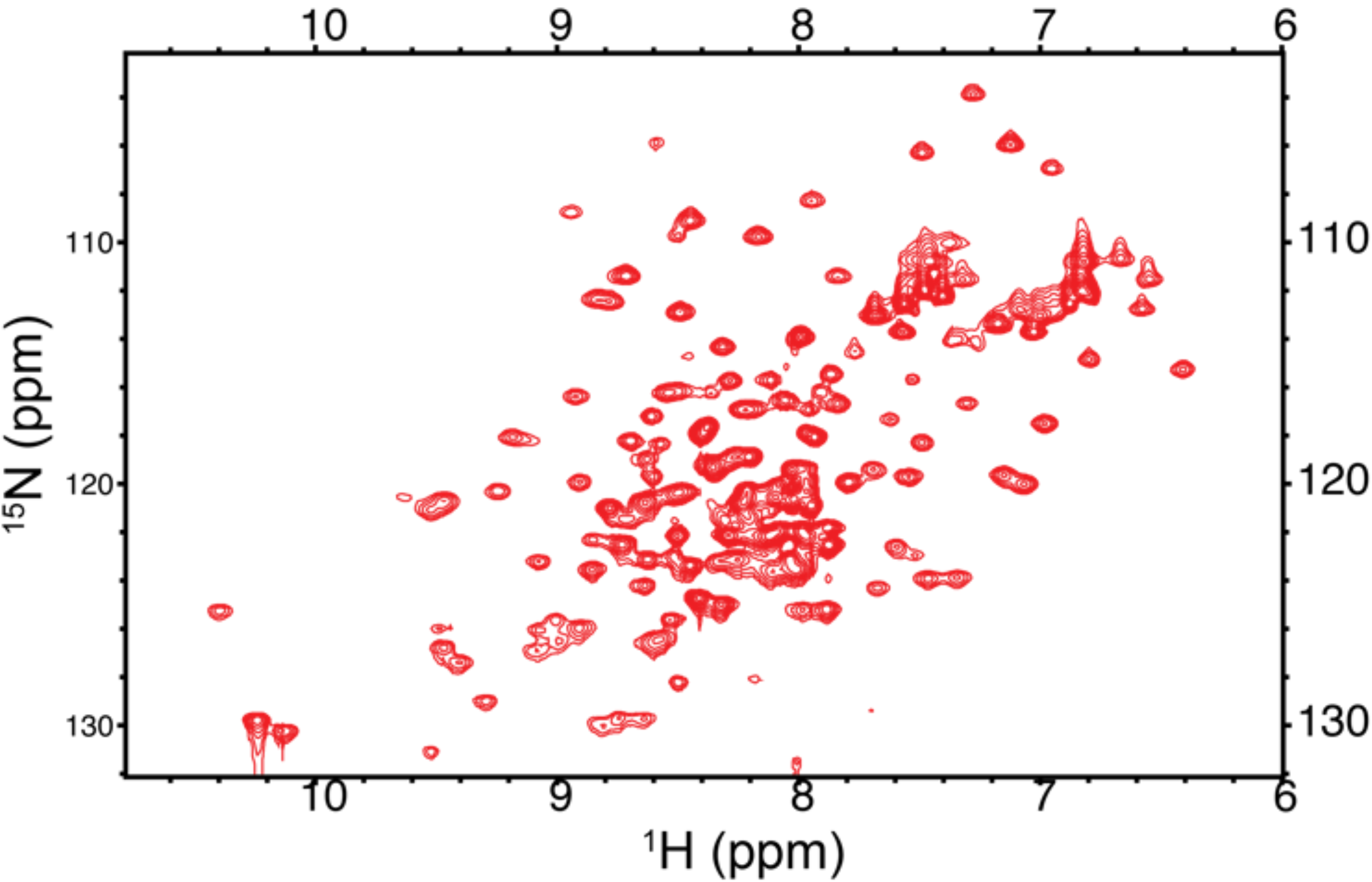
2D [^15^N, ^1^H]NMR spectrum of Sub16pro. 2D [^15^N, ^1^H] HSQC spectrum of Sub16pro (1 mM protein, 10 mM sodium phosphate pH7.4 and 100 mM NaCl) exhibits resonances with excellent chemical shift dispersion, indicative of a well-folded protein. The broad line-widths of resonances, however, indicate formation of higher order structure and intermediate conformational exchange.

### 3.5 Homology modeling and MD simulation studies of Sub16pro-catalytic domain complex

In order to investigate atomistic properties of Sub16pro that result in its structural flexibility, stand-alone homology models of Sub16pro or its cognate catalytic domain were generated. Human TPP1 (PDBID: 3EE6), with sequence similarity of 27.9% (e-value: e^-13^) and 34.2% (e-value: e^-54^) with Sub16pro and Sub16 catalytic domain, respectively, is the only available structure of any sedolisin family member with no available structure for independent stand-alone sedolisin pro domain. Although structures of stand-alone pro-domain of S8A family are available, they exhibit poor sequence similarity with Sub16pro. The homology models of Sub16pro and the catalytic partner were hence generated using equivalent domains in the structure of precursor human TPP1, as templates.

The conformational flexibility of the homology models was probed by MD simulation studies at 100 ns time scale. The Sub16 catalytic domain showed minor differences in C^α^ root mean-square deviation (RMSD) trajectories in initial 17-19 nanoseconds of simulations and eventually leading to a stable equilibrium throughout the rest of 100 ns of simulation. MD simulation studies with the stand-alone Sub16pro were more dramatic as it exhibited much larger structural changes as compared to the catalytic domain. An examination of the RMSD and radius of gyration (R_g_) for Sub16pro shows a general increase in both these measures across the simulations (Fig. 5A & B). The RMSD for Sub16pro increases to a high 7 Å and finally converging at 5.5 Å after 100 ns of simulations (Fig. 5A).

**Figure 5:**
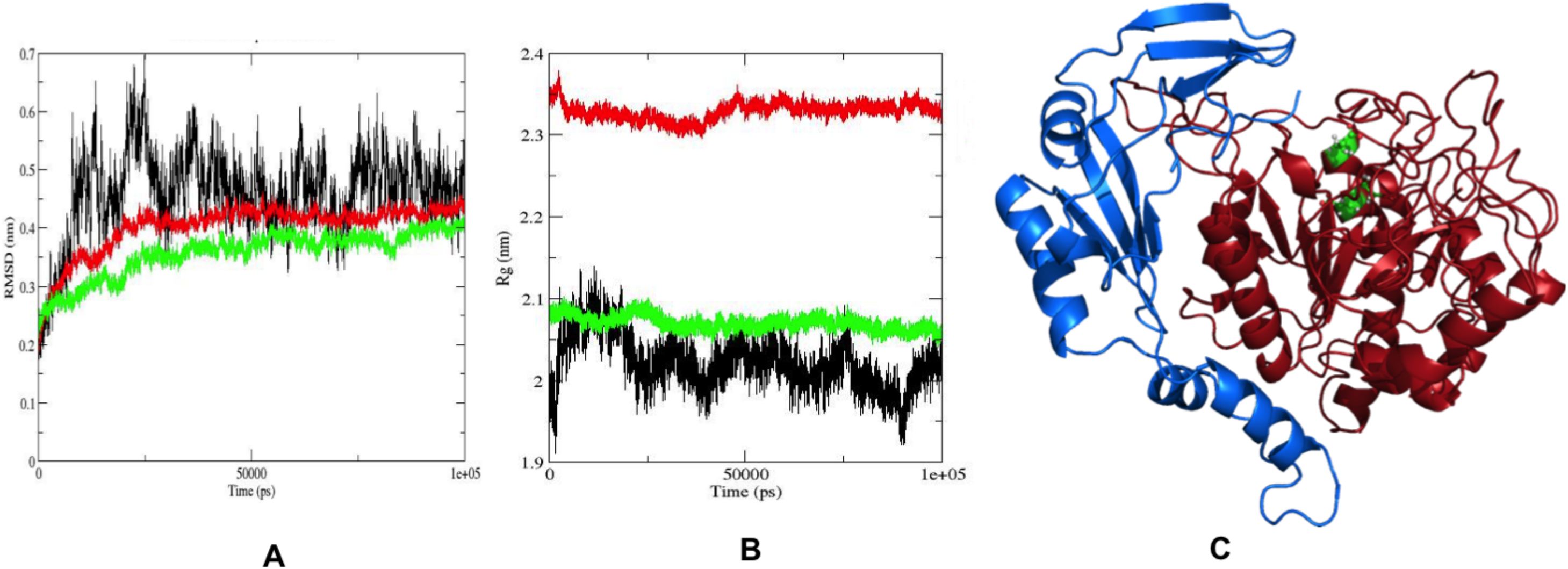
MD simulations on Sub16pro. (A) RMSD plots of Sub16pro (black), Sub16 catalytic domain (green) and Sub16pro-Sub16 catalytic domain complex (red) after 100ns of MD simulations (B) R_g_ plots of Sub16pro (black), Sub16 catalytic domain (green) and Sub16pro-Sub16 catalytic domain complex (red). (C) Three-dimensional structure of Sub16pro (blue)-Sub16 catalytic domain (brick red) complex after 100 ns of MD simulations. Catalytic triad is shown in green.

In order to identify a possible role of interactions between Sub16pro and its catalytic domain in stabilizing each other, a 100 ns MD simulation was next carried out between a complex of Sub16pro and the catalytic domain. The C^α^-RMSD plot for the complex is similar to that of the Sub16 catalytic domain, with minor differences in the trajectories in the initial 17-19 nanoseconds and eventually a stable equilibrium throughout the rest of 100 ns of simulation. A view of the Sub16pro-Sub16 catalytic domain, averaged over 100ns, shows a compact structure of the complex (Fig. 5C). The R_g_ for the complex, like that of the catalytic domain, again does not show much variations during the entire simulation run. In contrast, the R_g_-value of Sub16pro exhibits large fluctuations during the entire simulation run of 100 ns, suggesting large conformational dynamics during the MD run. Taken together, the investigation of conformational flexibility of Sub16pro, Sub16 catalytic domain and the Sub16pro-catalytic domain complex suggests that the standalone Sub16 pro domain has some inherent flexibility which is stabilized by catalytic domain, the detailed mechanism of which will require further investigations.

## 4. Conclusion

In conclusion, this is the first report of structural investigation on the prosegment of recently identified sedolisin family of subtilases in *T. rubrum*. Our results indicate that Sub16pro of *T. rubrum* may have inherent flexibility and conformational dynamics in the absence of catalytic domain that renders structure determination of ‘full length’ pro segment challenging. Secreted subtilases play an important role in the infective growth stages of *T. rubrum*. We identify a short subdomain of Sub16pro that might act as an inhibitor of catalytic domain of members of sedolisin family and aid in design of potential inhibitors. Furthermore, detailed structural characterization of pro segment with catalytic domain shall reveal information about mechanism of inhibition of active domain by the pro segment.

## Declaration of Competing interest

Authors declare no conflict of interest.

## Acknowledgements

We thank Dr. Shantanu Sengupta, CSIR-IGIB, India and the Mass spectrometry facility of CSIR-IGIB for help with mass spectrometry experiments; Dr. Deepak Nair, Regional Centre for Biotechnology, India (RCB) with diffraction data collection; the Proteomics Facility of RCB for N-terminal sequencing; CSIR-4PI for supercomputing facilities and Professor Stewart Shuman and Dr Krishna Murari Sinha for the pET28-His10-Smt3 plasmid. Senior Research Fellowship (SRF) to CL from CSIR, India, SRF to PA from DBT, India, SRF to RC from University Grants Commission, India and CSIR-Research Associateship to SHK from CSIR, India is gratefully acknowledged.

## Author contributions

CL: Conceptualization, Methodology, Investigation, Data Curation and Writing-original draft preparation. JB, PA,NSB,RC, SHK: Investigation, Methodology, Data Curation. BT: Conceptualization, Methodology, Data Curation, Resources, Supervision, Visualization, Writing-Review & Editing and Funding acquisition.

## Funding information

Data collection at ESRF (ID23, ID29) was facilitated by the ESRF Access Program of RCB which is supported by Department of Biotechnology (DBT), Government of India [Grant No.BT/INF/22/SP22660/2017]. DBT, India and ICGEB core fund for NMR facility at ICGEB, New Delhi is gratefully acknowledged. BT acknowledges support of DBT Project [No. BT/PR20790/MED/29/1130/2016] for work in his lab.

---

**Figure S1:**
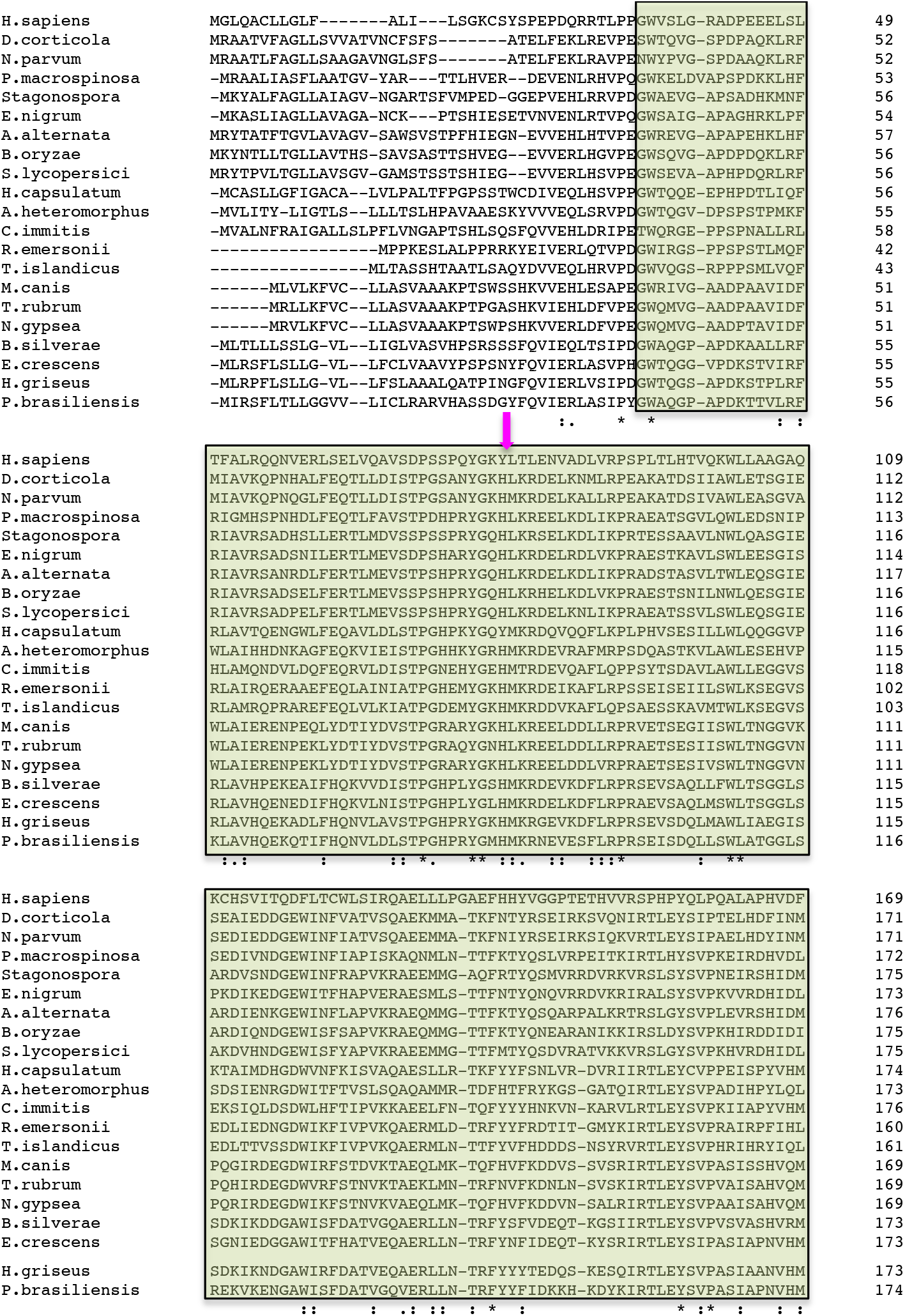
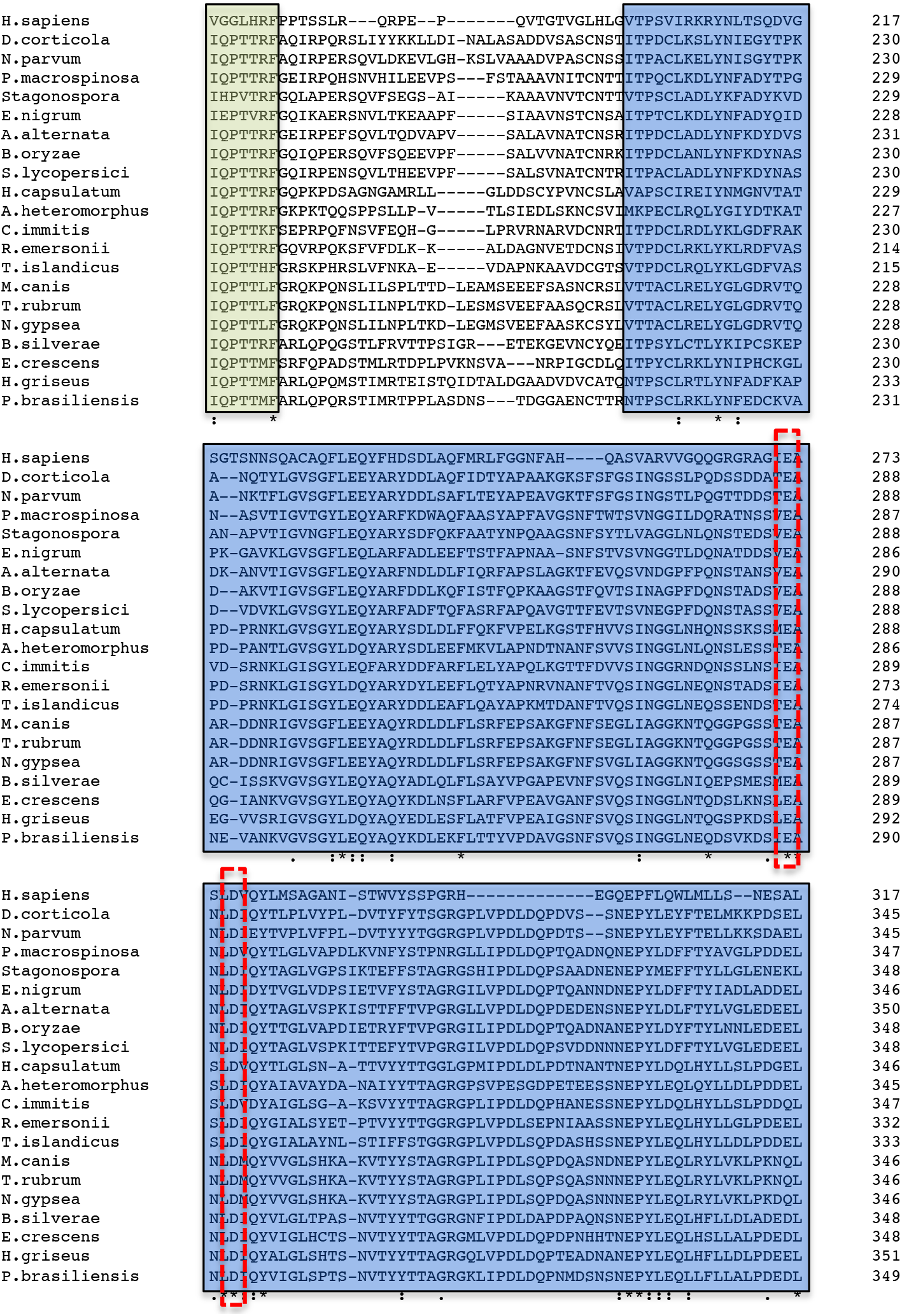
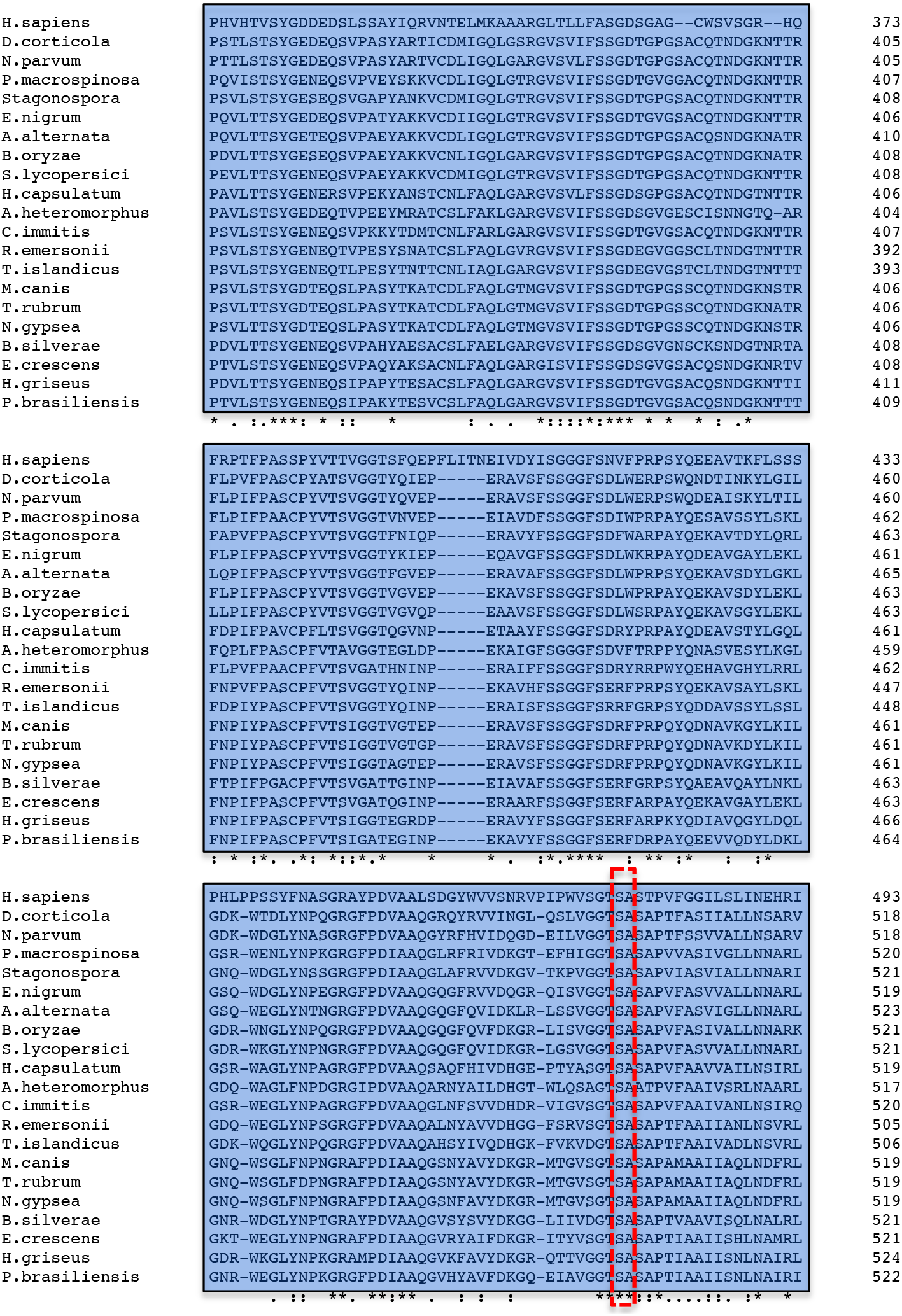
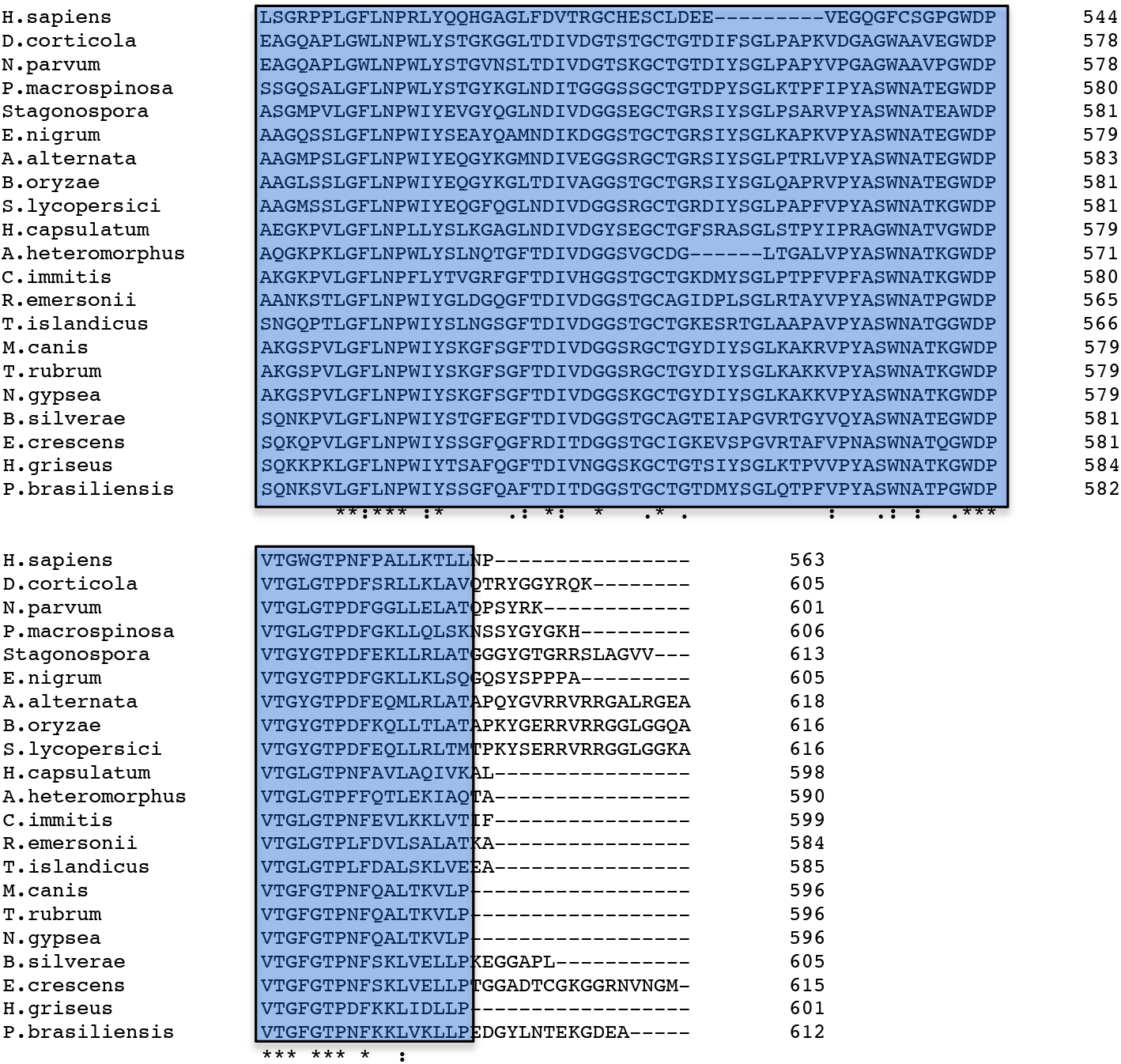
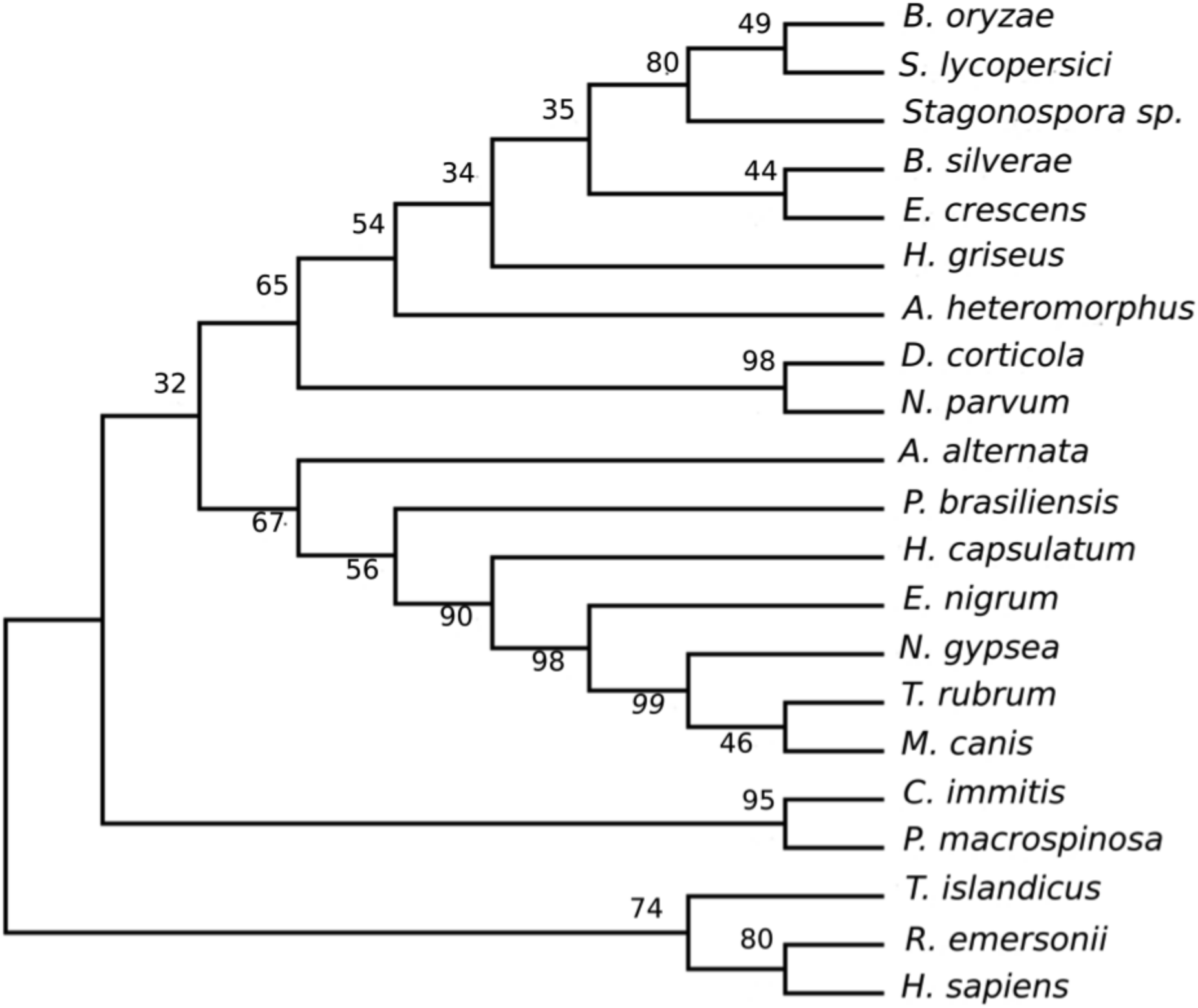
Sequence analysis of Sub16. (A) Sequence alignment of Sub16 (NCBI accession no.: KMQ48787, Locus_tag: HL42_0530) with selected sedolisin sequences of filamentous fungi. Pro domain is marked in yellow and catalytic domain is marked in blue. Catalytic triad is highlighted in a red box. Cleavage site of Sub16 pro domain is indicated by magenta arrow. (B) A phylogenetic tree was constructed by the Neighbor-Joining method in MEGA7. Bootstrap values are indicated. Amino acid sequence accession numbers are – *Alternaria alternata* (NCBI accession no.: XP_018388492.1), *Aspergillus heteromorphus* (NCBI accession no.: XP_025404410.1), *Bipolaris oryzae* (NCBI accession no.: XP_007684109.1), *Blastomyces silverae* (NCBI accession no.: KLJ12419.1), *Coccidioides immitis* (NCBI accession no.: XP_001244446.1), *Diplodia corticola* (NCBI accession no.: XP_020127950.1), *Emmonsia crescens* (NCBI accession no.: PGH35776.1), *Epicoccum nigrum* (NCBI accession no.: OSS54574.1), *Helicocarpus griseus* (NCBI accession no.: PGG95836.1), *Histoplasma capsulatum* (NCBI accession no.: EEH10792.1), *Homo sapiens* (NCBI accession no.: NP_000382.3), *Microsporum canis* (NCBI accession no.: XP_002850080.1), *Nannizzia gypsea* (NCBI accession no.: XP_003176669.1), *Neofusicoccum parvum* (NCBI accession no.: EOD49349.1), *Paracoccidioides brasiliensis* (NCBI accession no.: ODH48645.1), *Periconia macrospinosa* (NCBI accession no.: PVH98258.1), *Rasamsonia emersonii* (NCBI accession no.: XP_013326910.1), *Stagonospora* sp. SRC1lsM3a (NCBI accession no.: OAK99266.1), *Stemphylium lycopersici* (NCBI accession no.: KNG48502.1), *Trichophyton rubrum* (NCBI accession no.: KMQ48787) and *Talaromyces islandicus* (NCBI accession no.: CRG88197.1).

**Figure S2:**
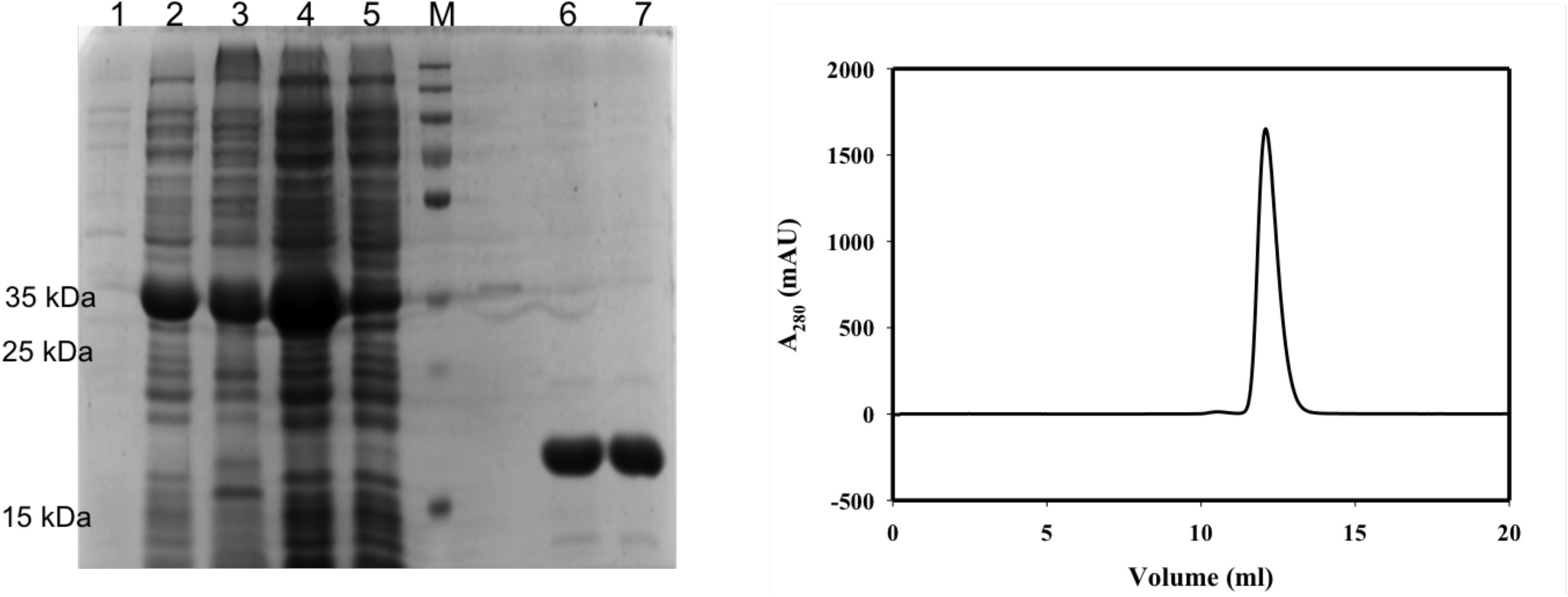
Purification of Sub16 pro. (A) SDS-PAGE of purification samples of Sub16pro. Lane 1: Uninduced control, Lane 2: Induced, Lane 3: Lysed Pellet, Lane 4: Supernatant, Lane5: Flow through (Ni-NTA column), Lane 6, 7: Elutes (from Ni-NTA column), Lane M: Molecular weight standards (in kDa) (B) Gel filtration chromatography of Sub16pro.

